# Global change differentially modulates coral physiology and suggests future shifts in Caribbean reef assemblages

**DOI:** 10.1101/2021.07.13.452173

**Authors:** Colleen B Bove, Sarah W Davies, Justin B Ries, James Umbanhowar, Bailey C Thomasson, Elizabeth B Farquhar, Jessica A McCoppin, Karl D Castillo

## Abstract

Global change driven by anthropogenic carbon emissions is altering ecosystems at unprecedented rates, especially coral reefs, whose symbiosis with algal endosymbionts ise particularly vulnerable to increasing ocean temperatures and altered carbonate chemistry. Here, we assess the physiological responses of the coral holobiont (animal host + algal symbiont) of three Caribbean coral species from two reef environments after exposure to simulated ocean warming (28, 31 °C), acidification (300 - 3290 μatm), and the combination of stressors for 93 days. We used multidimensional analyses to assess how multiple coral holobiont physiological parameters respond to ocean acidification and warming. Our results demonstrate significantly diminishing holobiont physiology in *S. siderea* and *P. astreoides* in response to projected ocean acidification, while future warming elicited severe declines in *P. strigosa*. Offshore *S. siderea* fragments exhibited higher physiological plasticity than inshore counterparts, suggesting that this offshore population has the capacity to modulate their physiology in response to changing conditions, but at a cost to the holobiont. Plasticity of *P. strigosa* and *P. astreoides* was not clearly different between natal reef environments, however, temperature evoked a greater plastic response in both species. Interestingly, while these species exhibit unique physiological responses to ocean acidification and warming, when data from all three species are modeled together, convergent stress responses to these conditions are observed, highlighting the overall sensitivities of tropical corals to these stressors. Our results demonstrate that while ocean warming is a severe acute stressor that will have dire consequences for coral reefs globally, chronic exposure to acidification may also impact coral physiology to a greater extent than previously assumed. The variety of responses to global change we observe across species will likely manifest in altered Caribbean reef assemblages in the future.

## Introduction

Human-induced global change is driving unprecedented variations in how global ecosystems function, from increases in terrestrial dryness (Greve et al., 2018) and severe storm activity across lower latitudes (Delworth et al., 2016), to altering species’ ranges globally (Habary et al., 2017; MacLean & Beissinger, 2017). Coral reefs are a prime example of an ecosystem heavily impacted by global change, particularly by ocean acidification and warming (Bove et al., 2021; Hoegh-Guldberg et al., 2007; Kleypas, 2019; Knowlton, 2001). Ocean acidification and warming are predicted to affect many marine ecosystems by reducing ecosystem complexity and function, especially for organisms with longer generational times and thus fewer opportunities to adapt to changing conditions (Nagelkerken & Connell, 2015). Therefore, understanding the diversity of responses of tropical reef-building corals at both the species- and population-levels is critical for predicting future impacts of global change.

Quantification of coral calcification rates (e.g., rate of skeletal production) is an informative and common tool used to assess overall coral health under stress in both field and laboratory experiments (Allemand et al., 2010; Comeau et al., 2013; Crook et al., 2013; Enochs et al., 2014; Kenkel et al., 2015; Ries et al., 2010). Understanding changes in calcification rates is critical in evaluating how coral reefs will respond to global change owing to the ecological importance of new reef production. Previous work quantifying coral calcification rates under global change stressors has demonstrated a diversity of growth responses under stress, including both maintained and suppressed calcification rates (Bove et al., 2019; Comeau et al., 2013; Okazaki et al., 2017). Corals that are better able to maintain calcification rates under stress may accomplish this at a cost to other metabolic processes (Cohen & Holcomb, 2009; Von Euw et al., 2017). Although calcification rates are valuable measures of the coral stress response, they do not provide insight into how the individual components of the coral holobiont (animal host and dinoflagellate symbionts) respond to projected global change stressors. To address this gap, many studies now quantify physiological responses of the coral host and algal symbionts under stress to investigate coral responses more thoroughly (Kenkel et al., 2015; Rodolfo-Metalpa et al., 2010; Schoepf et al., 2013).

Tropical reef-building corals depend on the maintenance of an endosymbiotic relationship with photosynthetic dinoflagellates (family Symbiodiniaceae) for a significant portion of their energetic needs (Muscatine et al., 1981). However, this relationship often breaks down under times of severe or prolonged stress, especially with increasing seawater temperatures, resulting in the phenomenon known as ‘coral bleaching’ (Anthony et al., 2008; Brown, 1997; Glynn, 1996). As corals bleach in response to ocean acidification and especially warming, physiological processes, including calcification and gametogenesis (Szmant & Gassman, 1990), deteriorate. Thus, as the symbiosis between the coral host and algal symbionts breaks down, both components of the holobiont are likely to exhibit closely integrated physiological responses. Indeed, previous work has observed that greater coral tissue biomass follows increased symbiont density and chlorophyll a content in several Caribbean reef-building coral species (Fitt et al., 2000), highlighting the intrinsic relationship with algal symbionts to support the coral host’s energy budget.

Coral tissue biomass and energy reserves (e.g., lipid, protein, carbohydrate) are important aspects of overall coral health (Rodrigues & Grottoli, 2007; Schoepf et al., 2013) that provide insight into resilience and recovery capacity in response to environmental stressors. Although energy reserves are extremely important in understanding the coral host response to stress, few studies have looked into how the combination of ocean acidification and warming influence these quantities (Schoepf et al., 2013; Towle et al., 2015). Coral tissue biomass relies on the equilibrium between energy sources and expenditures; thus, corals with already low biomass (i.e., low energy reserves) may experience heightened vulnerability under environmental stress (Thornhill et al., 2011) and may explain some of the variation of physiological responses to stress within and between species (Bove et al., 2019; Okazaki et al., 2017). However, studies have demonstrated that corals may not always consume energy reserves under environmental stress (Schoepf et al., 2013) or increase metabolic processes (Edmunds, 2012). Instead, corals may use other physiological mechanisms as coping tools to maintain growth and host energy reserves, such as relying more on algal symbionts whose photosynthesis is fertilized under conditions of elevated *p*CO_2_ (Guillermic et al., 2021).

To assess the physiological responses of Caribbean coral holobionts (animal host and algal symbionts) to independent and combined ocean acidification (300 - 3290 μatm) and warming (28, 31 °C), we conducted a 93-day common-garden experiment on 3 species of corals (*Siderastrea siderea, Pseudodiploria strigosa, Porites astreoides*) and quantified coral host energy reserves (total protein, carbohydrate, lipid) and algal symbiont physiology (cell density, chlorophyll a concentration, color intensity). Based on previous similar work, we hypothesized that (1) coral holobionts are more susceptible to thermal stress than acidification, (2) physiological responses are highly species-specific, (3) coral hosts are more susceptible than their algal symbionts, and (4) physiological plasticity dictates coral holobiont under global change. Our results highlight the diversity of physiological responses that Caribbean corals exhibit in response to projected global change, which will ultimately drive changes in the abundance and distribution of these species.

## Methods

### Experimental design

This study further investigates the physiological responses of the corals assessed in Bove et al (2019) thus, detailed descriptions of experimental setup can be found there. Briefly, six colonies each of three Caribbean reef-building corals (*Siderastrea siderea, Pseudodiploria strigosa, Porites astreoides*) were collected from inshore (Port Honduras Marine Reserve) and offshore (Sapodilla Cayes Marine Reserve) reef environments from the southern portion of the Belize Mesoamerican Barrier Reef System. Corals were immediately transported to Northeastern University’s Marine Science Center. Colonies were sectioned into eight equally-sized fragments and maintained in one of eight experimental treatments (three replicate tanks per treatment) for 93 days after a 23-day recovery time and a 20-day acclimation period. The eight treatments encompassed four *p*CO_2_ treatments corresponding to pre-industrial, current-day (*p*CO_2_ control), moderate end-of-century *p*CO_2_, and an extreme *p*CO_2_ level all crossed with two temperatures corresponding to the corals’ approximate present-day summer mean (28 °C) and projected end-of-century summer warming (31 °C). These *p*CO_2_-temperature combinations resulted in eight triplicate (24 tanks total) treatments: *p*CO_2_ (±SD) = 311 (±96), 405 (±91), 701 (±94), 3309 (±414) μatm at T (±SD) = 28°C (±0.4); and *p*CO_2_ (±SD) = 288 (±65), 447 (±152), 673 (±104), 3285 (±484) μatm at T (±SD) = 31.0°C (±0.4).

Experimental tanks were filled with filtered natural seawater with a salinity of 31.7 (±0.2) and were illuminated on a 10:14 light-dark cycle with photosynthetically active radiation of approximately 300 μmol photons m^−2^ s^−1^. Temperature, salinity and pH were measured every other day throughout the experiment and total alkalinity (TA) and dissolved inorganic carbon (DIC) were analyzed every 10 days with a VINDTA 3C (Marianda Corporation, Kiel, Germany). Temperature, salinity, TA and DIC were used to calculate carbonate parameters using CO_2_SYS (Pierrot et al., 2006) with Roy *et al*. (1993) carbonic acid constants K_1_ and K_2_, Mucci’s value for the stoichiometric aragonite solubility product (Mucci, 1983), and an atmospheric pressure of 1.015 atm. At the completion of the experimental period, corals were immediately flash-frozen in liquid nitrogen and transported to the University of North Carolina at Chapel Hill. Coral tissue was removed from the skeleton using seawater with an airbrush and stored in 50 mL conical tubes at −80°C until further processing.

### Host and symbiont physiological parameter assessments

Preserved coral holobiont tissue slurries were homogenized with a *Tissue-tearor* (BioSpec Products; Bartlesville, Oklahoma, USA) for several minutes and vortexed for 5 seconds, after which 1.0 mL of slurry was aliquoted for algal symbiont density analysis. Algal symbiont aliquots were dyed with 200 μL of a 1:1 Lugol’s iodine and formalin solution and cell densities were quantified by performing at least 3 replicate counts of 10 μL samples using a hemocytometer (1 x 1 mm; Hausser Scientific, Horsham, Pennsylvania, USA) and a compound microscope. Algal symbiont densities were standardized to total tissue volume and previously measured coral surface area (10^6^ cells per cm^2^) (Bove et al., 2019). Remaining tissue slurry was centrifuged at 4400 rpm for 3 minutes to separate the coral host and algal symbiont fractions, and the host fraction was poured off from the symbiont pellet. Chlorophyll a pigment was extracted from the algal pellet by adding 40 mL of 90% acetone to the conical tube at −20°C for 24 hours. Samples were diluted by adding 0.1 mL of extracted chlorophyll a sample to 1.9 mL of 90% acetone. If samples were too high or too low for detection on the fluorometer, samples were reanalysed by either diluting or concentrating the sample, respectively. Extracted chlorophyll a content was measured using a Turner Design 10-AU fluorometer with the acidification method (Parsons et al., 1984) and expressed as the μg of pigment per cm^2^ of coral tissue surface area.

Coral host supernatant was aliquoted (1 mL each) for total protein, carbohydrate, and lipid analysis, and stored at −80 °C. Glass beads were added to total protein aliquots, vortexed for 15 minutes, and centrifuged for 3 minutes at 4000 rpm. Duplicate samples were prepared with 235 μL of seawater, 15 μL of protein aliquot, and 250 μL of Bradford reagent (*Thermo Scientific*) and left for ca. 20 minutes. Coral host total protein samples were read at 562 nm on a spectrophotometer (Eppendorf BioSpectrometer® basic; Hamburg, Germany) in duplicates and were expressed as mg per cm^2^ coral tissue surface area. For coral host carbohydrate, 25 μL of phenol was added to 1000 μL of diluted coral host slurry and vortexed for 3 seconds before immediately adding 2.5 mL concentrated sulphuric acid (H_2_SO_4_). Samples were incubated at room temperature for 1 minute and then transferred to a room temperature water bath for 30 minutes (Masuko et al., 2005). Finally, 200 μL of each standard and sample was pipetted into a 96-well plate in triplicate and read on a spectrophotometer at 485 nm (BMG LABTECH POLARstart Omega; Cary, North Carolina, USA). Total carbohydrate was expressed as mg per cm^2^ coral tissue surface area (Bove & Baumann, 2021a). Coral host lipids were extracted following the Folch Method (Folch et al., 1957) by adding 600 μL of chloroform (CHCl_3_) and methanol (CH_3_OH) in a 2:1 ratio to 600 μL of host slurry and placed on a plate shaker for 20 minutes before adding 160 μL of 0.05M sodium chloride (NaCl). Tubes were inverted twice and then centrifuged at 3000 rpm for 5 minutes. Finally, the lipid layer was removed and 100 μL was pipetted in triplicate into a 96-well plate for colorimetric assay. The lipid assay was performed by adding 50 μL of CH_3_OH to each well before evaporating the solvent at 90 °C for 10 minutes. Next, 100 μL of H_2_SO_4_ was added to every well, incubated at 90 °C for 20 minutes, and cooled on ice for 2 minutes before transferring 75 μL of each sample into a new 96-well plate. Background absorbance of the new plate was read at 540 nm on a spectrophotometer before adding 34.5 μL of 0.2 mg/mL vanillin in 17% phosphoric acid to each well. The plate was read again at 540 nm and coral host lipid concentrations were normalised to coral surface area (mg per cm^2^) (Bove & Baumann, 2021b; Cheng et al., 2011).

Coral color intensity was analysed from images of every fragment with standardized color scales taken at every time point during the experiment. Color balance was adjusted using a custom Python script that took a square of pixels as a white standard (50 x 50) on each image to adjust the color balance until it was true white. The total red, green, blue, and sum of all color channel intensities were measured following (Winters et al., 2009) using the MATLAB macro “AnalyzeIntensity” for either 10 (*S. siderea* and *P. astreoides*) or 20 (*P. strigosa*) quadrats of 25 x 25 pixels on each coral fragment. The resulting values act as a measure of brightness, with higher brightness values correlating with pigment lightening (i.e., coral bleaching), thus, data were inverted so that lower values represent reduced coral pigmentation. The sum of all color channels (red, green, blue) resulted in a stronger correlation with symbiont physiology (chlorophyll a and cell density) in *S. siderea* and *P. strigosa*, while the red channel alone was best in *P. astreoides*.

### Coral holobiont physiology analyses

Principal component analysis (PCA) (function *prcomp*) of scaled and centered physiological parameters (host carbohydrate, host lipid, host protein, algal symbiont chlorophyll a, algal symbiont cell density, holobiont calcification rate as previously for the same samples in Bove et al (2019)) were employed to assess the relationship between physiological parameters and treatment conditions for each coral species. Main effects (temperature, *p*CO_2_, reef environment) were evaluated with PERMANOVA using the *adonis2* function (*vegan* package; version 2.5.7 (Oksanen et al., 2020)).

Correlations of all physiological parameters were assessed to determine the relationships between parameters within each species. The Pearson correlation coefficient (R^2^) of each comparison was calculated using the corrgram package (version 1.13 (Wright, 2018)) and the significance was calculated using the *cor.test* function. These relationships were then visualized through simple scatterplots.

Physiological plasticity of each experimental fragment was calculated for each species using all seven PCs calculated above as the distance between an experimental fragment and the control (420 μatm; 28°C) fragment from that same colony. The effects of treatment (*p*CO_2_ and temperature) and natal reef environment on calculated distances were assessed using generalized linear mixed effects models (function *lmer*) with a Gamma distribution and log-link and a random effect for colony. The best-fit model was selected as the model with the lowest AIC for each species (**Table S1**). Natal reef environment was only a significant predictor of plasticity in *S. siderea;* thus, samples were pooled across reef environments for both *P. strigosa* and *P. astreoides.* Parametric bootstraps were performed to model mean response and 95% confidence intervals with 1500 iterations and significant effects were defined as non-overlapping confidence intervals. Marginal and conditional R^2^ values of the best fit models were calculated using the *r2_nakagawa* function in the rcompanion package (version 2.4.1 (Mangiafico, 2021)). All figures and statistical analyses were carried out in R version 3.6.3 (R Core Team, 2018) and the accompanying data and code can be freely accessed on GitHub (github.com/seabove7/Bove_CoralPhysiology) and Zenodo (10.5281/zenodo.5093907).

## Results

### Principal component analysis

Two principal components (PCs) explained approximately 66% of the variance in physiological responses of the *S. siderea* holobiont to ocean acidification and warming treatments (**Figure 1A**). PC1 was driven by differences in algal symbiont physiology (chlorophyll a, cell density), while PC2 represented an inverse relationship between host energy reserves (lipid, protein, carbohydrate) and calcification rates and color intensities. Overall, lower *p*CO_2_ and temperature resulted in improved *S. siderea* holobiont physiology (**Figure 1A**). Treatment *p*CO_2_ predominantly drove *S. siderea* physiological responses (p < 0.001; **Table S2**), while temperature and reef environment were not as strong drivers of physiological responses (p < 0.01 and p < 0.01, respectively; **Table S2; Figure S1A**).

**Figure 1.**
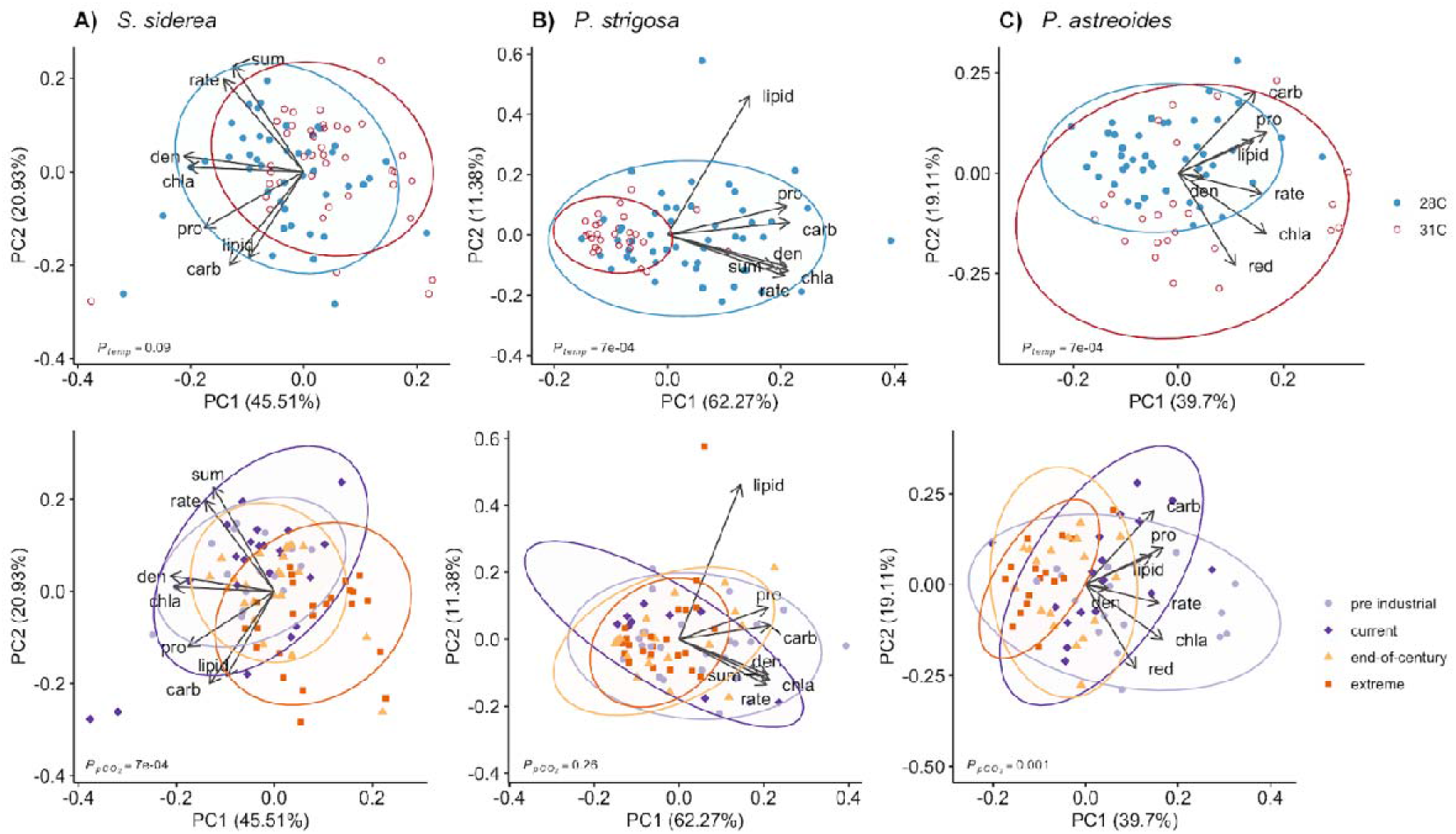
Principal component analysis (PCA) of all coral holobiont physiological parameters for (**A**) *S. siderea*, (**B**) *P. strigosa*, and (**C**) *P. astreoides* after 93 days of exposure to different temperature and *p*CO_2_ treatments. PCAs of each species are depicted by temperature treatment (28 °C blue; 31 °C red) in the top row and by *p*CO_2_ in the bottom row (pre industrial [300 μatm], light purple; current day [420 μatm], dark purple; end-of-century [680 μatm], light orange; extreme [3290 μatm], dark orange). Arrows represent significant (p < 0.05) correlation vectors for physiological parameters and ellipses represent 95% confidence based on multivariate t-distributions.

For *P. strigosa*, 74% of the variance in the holobiont responses to treatment was explained by two PCs (**Figure 1B**). PC1 explained most of the variation of physiological parameters with the exception of host lipid content, which was represented in PC2. Holobiont physiology of *P. strigosa* was clearly diminished under warming and was generally higher in the lower *p*CO_2_ treatments (**Figure 1B**). Treatment temperature (p < 0.001; **Table S2**), *p*CO_2_ (p < 0.01; **Table S2**), and natal reef environment all varied significantly with coral holobiont physiology (p < 0.001; **Table S2; Figure S1B**).

For *Porites astreoides,* the first two PCs explained about 59% of the total variance in holobiont response to treatment (**Figure 1C**). Samples separated most clearly along PC1 driven primarily by calcification rate and algal symbiont density, while PC2 exhibited an inverse relationship between host total carbohydrate and color intensity. Overall, lower *p*CO_2_ drove improved *P. astreoides* holobiont physiology, while elevated temperature resulted in greater holobiont physiology (**Figure 1C**). Temperature (p < 0.001; **Table S2**) and *p*CO_2_ (p < 0.001; **Table S2**) drove separations in *P. astreoides* holobiont physiology, while reef environment was not significant (p = 0.82; **Table S2; Figure S1C**).

### Correlations of physiological parameters

Coral holobiont physiological parameters were generally positively correlated with one another within each of the three species. Correlations between *S. siderea* holobiont physiological parameters identified 15 significant relationships out of all 21 possible comparisons (**Figure 2A**). Of those significant correlations, six resulted in a Pearson’s correlation coefficient (R^2^) equal to or greater than 0.5, with the strongest relationship identified between symbiont density and chlorophyll a (R^2^ = 0.72).

**Figure 2.**
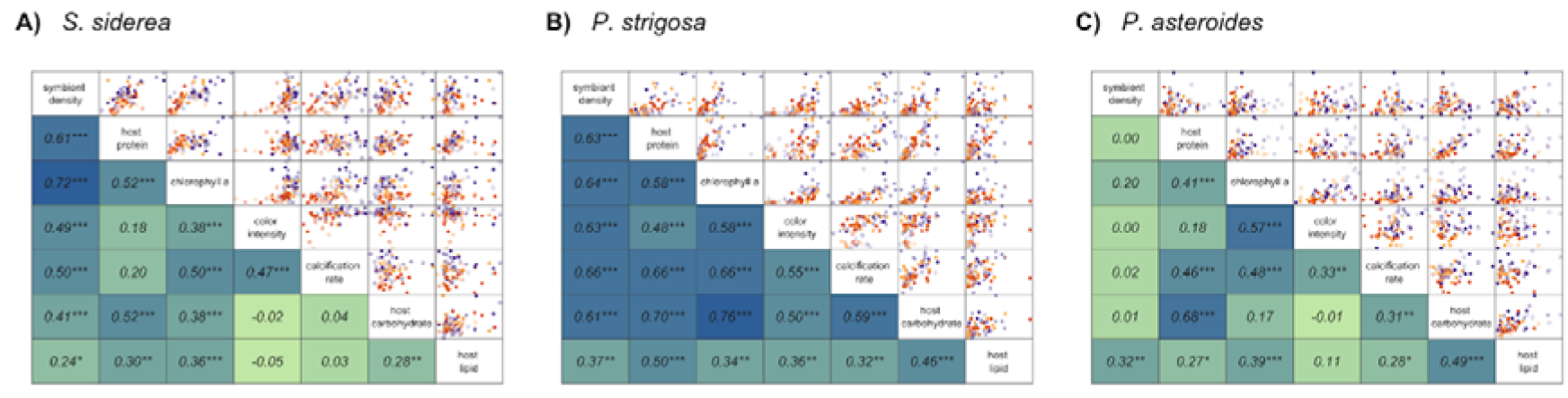
Coral holobiont physiological parameter scatter plots (top) and correlation matrices (bottom) for (**A**) *S. siderea*, (**B**) *P. strigosa*, and (**C**) *P. astreoides* showing pairwise comparisons of within each species. Scatter plots of each pairwise combination of physiological parameters are displayed on the top with temperature treatment depicted by shape (28 °C closed points; 31 °C open points) and *p*CO_2_ treatment depicted by color (pre industrial [300 μatm], light purple; current day [420 μatm], dark purple; end-of-century [680 μatm], light orange; extreme [3290 μatm], dark orange). Strengths of the correlations (R^2^ via Pearson correlation coefficients) between each pairwise combination of physiological parameters are indicated by darker shades of blue on the bottom with significance depicted by asterisks according to significance level (* p < 0.05; ** p < 0.01; *** p < 0.001). R^2^ and significance levels correspond to the scatter plot at the intersection between two physiological parameters.

All pairwise physiological parameters were significantly correlated with one another in *P. strigosa,* and, of those, 15 correlations exhibit moderate (R^2^ > 0.50) positive relationships (**Figure 2B**). Notably, the two strongest correlations were host carbohydrate vs. host protein (R^2^ = 0.70) and host carbohydrate vs. chlorophyll a (R^2^ = 0.76).

Compared to both *S. siderea* and *P. strigosa*, fewer physiological traits were significantly (p < 0.05) correlated with one another in *P. astreoides* (12 significant out of 21 total comparisons; **Figure 2C**). Of the significant correlations, only two pairwise comparisons resulted in a Pearson’s correlation coefficient greater than 0.5: chlorophyll a vs. color intensity (R^2^ = 0.57) and host carbohydrate vs. host protein (R^2^ = 0.68).

### Coral holobiont physiological plasticity

Physiological plasticity of offshore *S. siderea* fragments exhibited a positive linear trend with increasing *p*CO_2_ while the inshore fragments appear to respond in a parabolic pattern to *p*CO_2_, with the lowest calculated distances occurring at 420 μatm, 31 °C and 680 μatm (both temperatures) (**Figure 3A**). Further, offshore *S. siderea* fragments exhibited higher plasticity in the extreme *p*CO_2_ treatment than in inshore fragments reared in the pre-industrial, current-day, and extreme *p*CO_2_ treatments, regardless of temperature (**Figure 3A; Table S3**).

**Figure 3.**
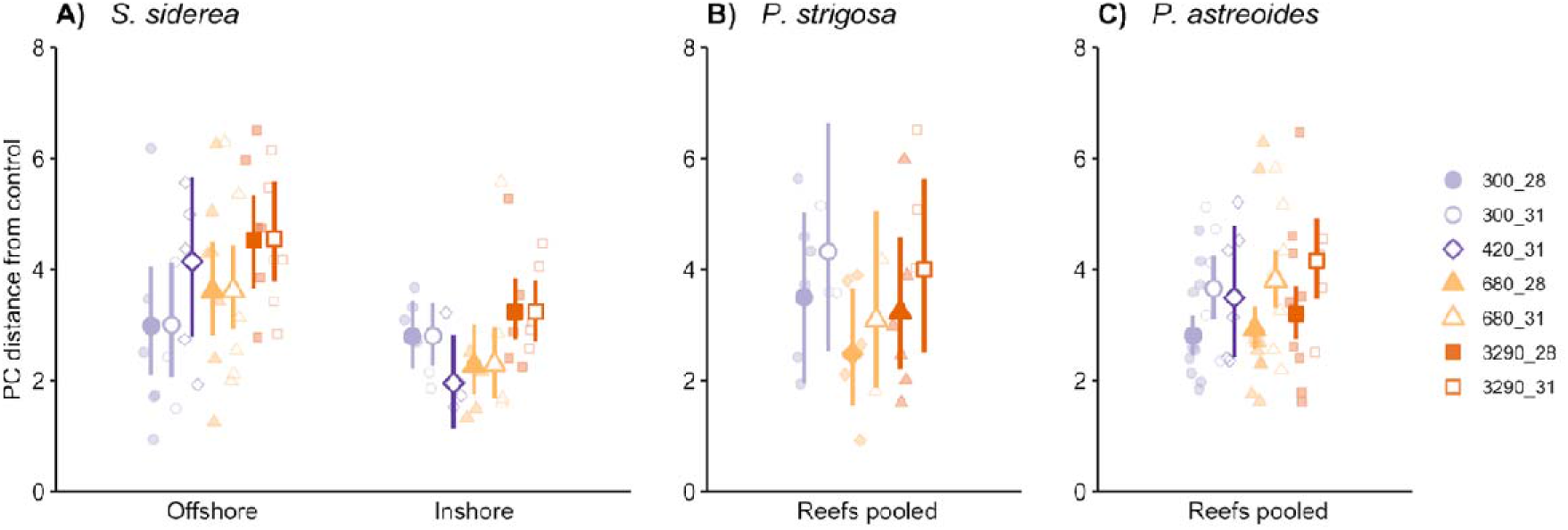
Holobiont physiological plasticity of (**A**) *S. siderea*, (**B**) *P. strigosa*, and (**C**) *P. astreoides* after 93-day exposure to experimental treatments. Higher values represent greater plasticity in coral holobiont samples. Natal reef environment is depicted along the x axis for *S. siderea*, however, *P. strigosa* and *P. astreoides* samples were pooled by reef environment. *p*CO_2_ treatment is depicted by color and shape (pre industrial [300 μatm], light purple; current day [420 μatm], dark purple; end-of-century [680 μatm], light orange; extreme [3290 μatm], dark orange) and temperature is represented as either closed (28 °C) or open (31 °C) symbols. Symbols and bars indicate modeled means and 95% confidence intervals.

Plasticity of *P. strigosa* and *P. astreoides* was not clearly different between colonies based on natal reef environments (see **Table S1**). No clear differences in physiological plasticity in response to treatment were identified in *P. strigosa* (**Figure 3B; Table S3**), however, this is likely due to reduced sample sizes in this analysis as a result of only five colonies (N_offshore_ = 3, N_inshore_ = 2) present in the control treatment for distance calculations.

Elevated temperatures generally resulted in higher plasticity of *P. astreoides* compared to control temperatures (**Figure 3C; Table S3**), however, this trend was not clearly different within each *p*CO_2_ treatment. Physiological plasticity of *P. astreoides* was significantly lower in both the pre-industrial and end-of-century *p*CO_2_ treatments at control temperatures than that measured in the extreme *p*CO_2_ treatment combined with the elevated temperature.

### Species differences in coral holobiont physiology

The first two PCs of the combined holobiont physiology explained about 62% of the total variance across samples (**Figure 4**). In general, fragments of *S. siderea* contained higher chlorophyll a content, host carbohydrate, and host lipid content, while *P. strigosa* fragments typically had greater host protein content accompanied by higher calcification rates, and fragments of *P. astreoides* were differentiated by their high symbiont densities (**Figure 4A; Table S5**). Despite being different coral species, coral holobiont physiology exhibited similar physiological responses to *p*CO_2_ and temperature treatments (**Figure 4B, 4C; Table S5**). As *p*CO_2_ or temperature increased, coral holobiont physiology was more constrained and exhibited convergent physiological responses under stress. Furthermore, corals from the inshore reef environment exhibited more constrained physiology than their offshore counterparts (**Figure S7; Table S5**).

**Figure 4.**
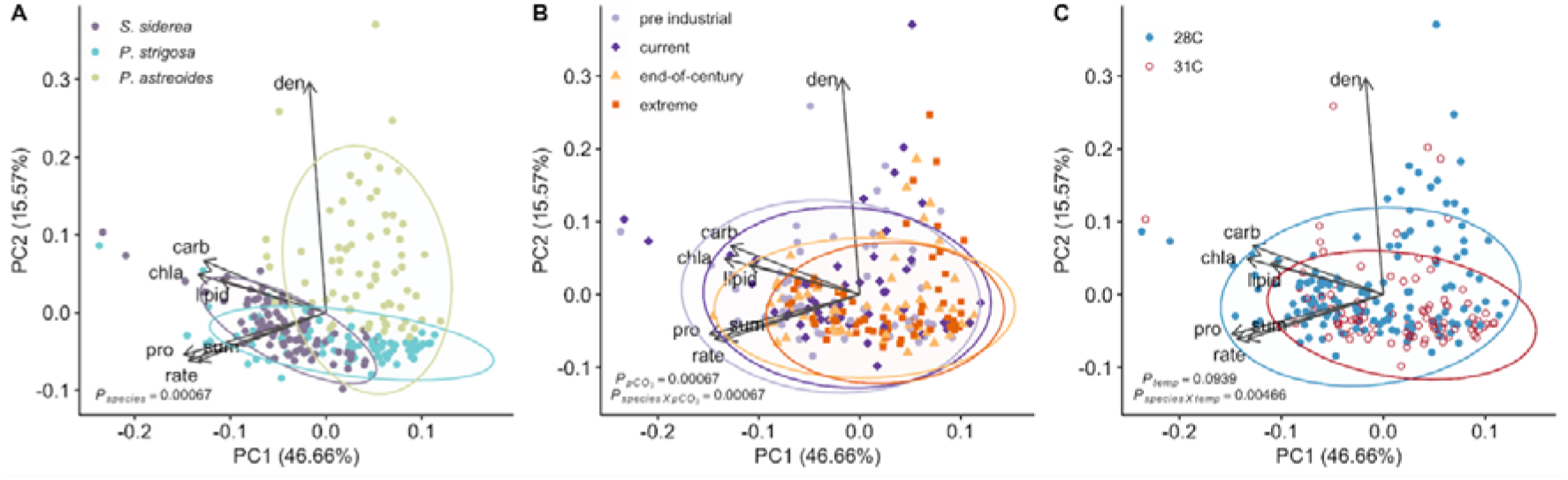
Principal component analysis (PCA) comparing the coral holobiont of all three species at the end of the experiment with samples clustered by (**A**) species, (**B**) *p*CO_2_ treatment, and (**C**) temperature treatment. Arrows represent significant (p < 0.05) correlation vectors for physiological parameters and ellipses represent 95% confidence based on multivariate t-distributions.

## Discussion

### Holobiont physiology highlights sensitivity of Caribbean corals to global change

Ocean acidification, warming, and the combination of the two stressors are expected to considerably reduce coral abundance throughout the Greater Caribbean by the end of this century (Cornwall et al., 2021). We demonstrate a variety of coral holobiont responses to simulated ocean acidification and warming scenarios that provide insight into how multiple coral species may respond to global change. Understanding the individual physiological responses of the coral host and their algal symbionts provides valuable insight into the relationship between partners within the coral holobiont. However, to better predict how coral holobionts will respond to global change, it is necessary to assess how the physiological parameters of both partners will respond. For example, we found that *p*CO_2_ treatment clearly drove differences in the coral holobiont physiology of both *S. siderea* and *P. astreoides* (**Figure 1**), however, these effects were not clear when assessing individual physiological parameters on their own within a species (**Figure S6**). Indeed, several previous studies have reported mixed holobiont physiological responses to elevated *p*CO_2_, from no difference in coral host energy reserves (Schoepf et al., 2013) to reduced symbiont density and productivity loss (Anthony et al., 2008; Edmunds, 2012). These effects of *p*CO_2_ highlight the complexity of the responses of coral holobionts under stress (Hoadley et al., 2019; Schoepf et al., 2013; Weis, 2010) and suggest that, by limiting assessments to only a few physiological parameters, studies may miss important changes to the coral holobiont.

Coral holobiont physiologies of all three species were also modulated by temperature, although these impacts were more variable. *Siderastrea siderea* and *P. strigosa* both exhibited declines in holobiont physiology under elevated temperature (31 °C) (**Figures 1, S6**), however, these declines were more pronounced in *P. strigosa,* especially through time (**Figures 1B, S6, S7**). Indeed, *P. strigosa* is known to be a more thermally sensitive coral species (Aichelman et al., 2021; Rippe et al., 2018; Scheufen et al., 2017) and this response is likely representative of the overall holobiont deterioration in response to thermal stress, which may lead to mortality under chronic exposure. Thermal events on coral reefs are generally considered more acute stress events (on the scale of hours to weeks) (Hughes et al., 2018). Thus, exposure of these corals to more than 90 days of constant elevated temperature may have elicited a more severe response in *P. strigosa* as is seen during mass bleaching events *in situ* for this species (Eakin et al., 2010). Conversely, elevated temperatures corresponded with improved holobiont physiological parameters in *P. astreoides.* These differences in coral holobiont thermal responses are not surprising given that *P. astreoides* is generally considered a more opportunistic coral that can persist in less-desirable conditions, including elevated temperatures (Darling et al., 2012; Green et al., 2008; Manzello et al., 2015). Conversely, *S. siderea* and *P. strigosa* are classified as ‘stress tolerant’ species with varying levels of susceptibility and resilience to environmental stress (Alemu & Clement, 2014; Neal et al., 2017; Okazaki et al., 2017; Venti et al., 2014). Despite some similarities in responses to ocean acidification and warming observed here, the different relationships between physiological parameters within each species likely interact to produce species-specific responses.

### Global change and species-specific drivers of physiological plasticity

On shorter ecological time scales – like those employed in this experiment – plasticity may be a coral’s most efficient response to global change, as it permits individual-level adaptation to a rapidly changing environment within a generation (Chevin et al., 2010; Fox et al., 2019). Plasticity has been identified as an important mechanism in coping with elevated *p*CO_2_ conditions in tropical corals (Camp et al., 2018; Putnam et al., 2016; Tambutté et al., 2015) and may predict how these organisms will perform under global change. Here, we assessed the physiological plasticity of the coral holobiont under elevated temperature and *p*CO_2_, and compared these responses across two natal reef environments (inshore vs. offshore). We found that *S. siderea* fragments from the offshore exhibited higher plasticity in response to extreme *p*CO_2_ (3290 μatm) compared to the inshore counterparts, and that the pattern of responses to increasing *p*CO_2_ differed between the two habitats (**Figure 3A; Table S3**). Our results suggest that offshore *S. siderea* fragments modulated their physiology to a greater extent than the inshore corals and may possess the physiological capacity to better persist under future ocean acidification. This higher plasticity, however, may come at a great physiological cost to the holobiont in the form of reduced fitness under optimum conditions (Chevin et al., 2010; DeWitt et al., 1998). Indeed, a reciprocal transplant experiment in southern Belize identified higher plasticity of offshore colonies of *S. siderea* compared to those from a nearshore environment (Baumann et al., 2021). The offshore colonies grew at a much higher rate when transplanted to the nearshore environment than in their natal environment (generally considered more ideal conditions) (Baumann et al., 2021), suggesting that plasticity in these corals may indeed come at the cost of growth in home or more ideal conditions.

Varying levels of plasticity in both *P. strigosa* and *P. astreoides* based on habitat have also been previously reported (Baumann et al., 2021; Kenkel & Matz, 2016), however, similar natal reef effects were not evident in either species in the current study (**Figure 3B-C; Table S3**). The small sample size of *P. strigosa* from both reef environments in this analysis likely contributed to the lack of plasticity differences between habitats for this species, while different measures of plasticity – physiological plasticity (present study) vs. gene expression plasticity (Kenkel & Matz, 2016) – may identify inconsistent plastic responses within *P. astreoides*. While neither species exhibited differential plasticity based on natal reef environment, both *P. strigosa* and *P. astreoides* appear to exhibit higher plasticity at the elevated temperature, though this is only statistically significant in *P. astreoides* (**Figure 3B-C; Table S3**). Interestingly, the higher plasticity associated with elevated temperatures in *P. strigosa* resulted in deterioration of holobiont physiological parameters, while higher plasticity in *P. astreoides* manifested as improved holobiont physiology (**Figure 1B-C**). These differences highlight how plasticity may result from physiological trade-offs in response to environmental change in some organisms (i.e., *P. strigosa*) (Chevin et al., 2010; DeWitt et al., 1998), and other organisms (i.e., *P. astreoides*) may benefit from such plastic responses to optimize physiology (Seebacher et al., 2015). Either way, it is clear that the role of plasticity in coral holobiont responses to global change is complex and merits further investigation to better understand species-specific levels of resilience.

Another explanation for varying susceptibilities across coral species under global change may relate to how physiological parameters are correlated to one another within the holobiont. For example, all holobiont physiological parameters were significantly correlated with one another for *P. strigosa* (**Figure 2B**), while only some correlations were significant for both *S. siderea* and *P. astreoides* (**Figure 2**). Notably, while symbiont density was significantly correlated with all holobiont parameters in *P. strigosa*, it was least correlated with host lipid content, which was best correlated with both host protein and carbohydrate (**Figure 2B**). This pattern suggests *P. strigosa* are consuming carbohydrate and protein stores in response to reduced symbiont density and photosynthetic efficiency, while lipid stores remain relatively unaltered, in line with previous work on coral energetics (Anthony et al., 2002; Gnaiger & Bitterlich, 1984). *Siderastrea siderea* exhibited similar relationships between symbiont density and all other physiological parameters; however, calcification rates were clearly more dependent on algal symbiont status than host energy reserves (**Figure 2A**). Interestingly, *P. astreoides* symbiont density only resulted in a significant correlation with lipid content, while chlorophyll a was a better predictor of most physiological parameters (**Figure 2C**). In fact, chlorophyll a and symbiont density resulted in one of the strongest correlations in both *S. siderea* and *P. strigosa*, while these two parameters were not significantly correlated in *P. astreoides*. This suggests that *S. siderea* and *P. strigosa* both rely on greater concentrations of algal symbionts with higher chlorophyll a content for autotrophically-derived carbon to support the coral host (Anthony & Fabricius, 2000; Muscatine et al., 1981), while *P. asteroides* is dependent on more efficient symbionts alone (Jones, 1997; Taylor, 1991). Additionally, the three species are known to host varying algal symbiont communities (e.g., *Siderastrea siderea* predominantly hosts *Cladocopium*; *P. strigosa* hosts *Cladocopium* and *Breviolum*; *P. astreoides* hosts *Breviolum* and *Symbiodinium* (LaJeunesse, 2002; LaJeunesse et al., 2018)) that may determine differing carbon allocation to the host as well as different thermal tolerances of the holobiont (Grégoire et al., 2017; Suggett et al., 2008). Although profiling of the algal symbiont community was outside the scope of the current study, both temperature and *p*CO_2_ can modulate the symbiosis between coral hosts and algal symbionts (Anthony et al., 2008; Baird et al., 2009; Brown, 1997; Coles & Brown, 2003). Therefore, given that algal symbiont community and physiology play a significant role in holobiont responses to global change stressors, these types of data should be obtained in future experiments to better understand differences between and within tropical coral species.

Interestingly, when comparing PCAs of host only (lipid, carbohydrate, protein) and symbiont only (chlorophyll a, symbiont density, color intensity) physiological parameters for each species, we found that algal symbionts were generally more impacted than the host by increasing *p*CO_2_ (**Figures S2-S5; Tables S5, S6**). For example, variance in *S. siderea* host physiology was not significantly explained by *p*CO_2_; however, *p*CO_2_ clearly affected symbiont physiology. This result suggests that coral bleaching was occurring under ocean acidification, but this bleaching did not affect host energy reserves (**Figure S2**). This pattern contrasts previous work demonstrating no declines in symbiont physiology under increased *p*CO_2_ (Crawley et al., 2010; Hii et al., 2009; Schoepf et al., 2013) and others highlighting greater transcriptomic plasticity of coral hosts in response to increasing *p*CO_2_ relative to their algal symbionts (Davies et al., 2018). Davies et al., (2018) interpreted this result as the coral host responding poorly to *p*CO_2_ stress. However, our results suggest that coral hosts were able to maintain energy reserves despite reductions in symbiont density and efficiency. There is debate on the exact relationship between the coral host and algal symbionts (i.e., mutualism vs. parasitism) as well as their relative roles in coral bleaching (A. C. Baker, 2001; Cunning & Baker, 2020; Wooldridge, 2010). While this symbiotic relationship was traditionally considered a mutualism, recent work has highlighted that this relationship is context dependent and, under specific circumstances, the algal symbionts may become more parasitic (D. M. Baker et al., 2018). Regardless, it is clear that understanding the varied responses of the different coral holobiont members is critical for predicting the future of tropical coral reefs.

### Global change drives similar physiological responses in Caribbean corals

Our results indicate species-specific relationships between physiological parameters within a coral holobiont that dictate responses to global change stressors and these patterns may separate the ‘winners’ from ‘losers’ (Fabricius et al., 2011; van Woesik et al., 2011). Comparisons across all experimental coral fragments highlight that *S. siderea* samples were largely differentiated by their higher host carbohydrate, host lipid, and chlorophyll a content, while *P. strigosa* fragments were associated with host protein and net calcification rates, and *P. astreoides* hosted the highest algal symbiont densities (**Figure 4A**). These physiological differences across species likely correspond to species-specific responses observed in this study and previous work assessing global change on tropical reef-building corals (Bove et al., 2019; Castillo et al., 2014; Kenkel et al., 2013; Okazaki et al., 2017), as well as patterns of resilience observed *in situ* (Green et al., 2008; Neal et al., 2017). For example, *S. siderea* has generally been considered a more resilient species in terms of survival and growth when reared under ocean acidification and warming conditions (Bove et al., 2019; Castillo et al., 2014; Okazaki et al., 2017). This resilience may be associated with this species’ maintenance of higher host carbohydrate reserves as a result of greater chlorophyll a content (Burriesci et al., 2012) along with increased host lipids reserves for long-term performance (Anthony et al., 2002; Gnaiger & Bitterlich, 1984). The association of proteins with *P. strigosa* is also noteworthy given that corals generally obtain proteins from their algal symbionts (Young et al., 1971). However, *P. strigosa* was the most bleached of the three species (see **Figures S6, S7**), suggesting that this species exhibited the largest variation in protein as a result of the loss of productive symbionts with warming. These differences across species not only highlight differences in the underlying physiology of Caribbean coral species, but may also assist in predicting responses to environmental stress.

Although the coral species examined here exhibit differing host and symbiont physiological responses, patterns of coral holobiont physiology converge under elevated temperature and *p*CO_2_, regardless of species (**Figure 4B, 4C**). As temperature and/or *p*CO_2_ increase, all three species exhibit similar declines in holobiont physiology. This pattern cautions that the broad classification of coral species as ‘resistant’ or ‘susceptible’ to environmental stressors based on individual physiological responses (Bove et al., 2019; Darling et al., 2012; Okazaki et al., 2017; Schoepf et al., 2013; Wall et al., 2017) overgeneralize sensitivity to future reef projections (Cornwall et al., 2021; Hoegh-Guldberg et al., 2007; Okazaki et al., 2017). The susceptibility observed here across all species is indicative of future Caribbean coral reef assemblages composed only of the most tolerant individuals within a species, despite some species level resilience to global change stressors.

## Conclusions

As global change continues, it is critical to understand species-specific coral holobiont responses to ocean acidification and warming scenarios in order to predict the future of Caribbean coral reef assemblages. Our results suggest that *S. siderea* may dominate reefs across the Caribbean due to its maintenance of tissue energy reserves and relatively unaltered symbiosis with their algal symbionts. Conversely, *P. strigosa* was unable to maintain any holobiont physiological parameters under warming, suggesting that this species is particularly vulnerable to thermal stress that will result in widespread bleaching and mortality. Finally, *P. astreoides* exhibited improved holobiont physiology under warming while ocean acidification drove reductions in the same physiological parameters, indicating that this species may also fare better than most on future coral reefs. Although these species had variable responses under these global change scenarios, all three exhibited overall deterioration in holobiont physiological parameters under the effects of ocean warming and acidification. Our results underscore the intricacies of coral holobiont physiology, both within and across species, in response to their environment and contribute to our understanding of the many ways that global change affects tropical coral reefs.

## Acknowledgements

We thank Belize Fisheries Department for all associated permits, the Toledo Institute for Development and Environment (TIDE) and the Southern Environmental Association (SEA) for their support. We also thank S. Patel, S. Swinea, F. Buckthal, J. Townsend, J. Boulton, and C. Lopazanski for assisting with preparing corals for physiological assays and the Marchetti, Septer, and Waters labs at UNC Chapel Hill for equipment and lab space use. This research was partially supported by the Women Diver Hall of Fame Sea of Change Foundation Marine Conservation Scholarship and Lerner-Gray Memorial Fund of the American Museum of Natural History Grants for Marine Research awarded to CBB. JBR acknowledges support from NSF BIO-OCE award #1437371.

## Data Accessibility

All data and code utilized in this manuscript are archived at Zenodo (10.5281/zenodo.5093907) and can be freely accessed on GitHub (github.com/seabove7/Bove_CoralPhysiology). Protocols for host carbohydrate and lipid assays can be accessed on protocols.io (carbohydrate: dx.doi.org/10.17504/protocols.io.bvb9n2r6; lipid: dx.doi.org/10.17504/protocols.io.bvcfn2tn).

