## Supplemental materials for "Global change differentially modulates coral physiology and suggests future shifts in Caribbean reef assemblages"

### **Supplemental Materials for manuscript:** *Global change differentially modulates coral physiology and suggests future shifts in Caribbean reef assemblages*


Colleen B Bove[1,2*], Sarah W Davies[1], Justin B Ries[3], James Umbanhowar[2,4], Bailey C Thomasson[4,5], Elizabeth B Farquhar[2,6], Jessica A McCoppin[4], Karl D Castillo[2,7]

[1] The Department of Biology, Boston University, Boston, Massachusetts, USA
[2] Environment, Ecology, and Energy Program, The University of North Carolina at Chapel Hill, Chapel Hill, North Carolina, USA
[3] Department of Marine and Environmental Sciences, Northeastern University, Nahant, MA, USA
[4] The Department of Biology, The University of North Carolina at Chapel Hill, Chapel Hill, North Carolina, USA
[5] Florida Fish and Wildlife Conservation Commission, St. Petersburg, Florida, USA
[6] Center for Marine Science, University of North Carolina Wilmington, Wilmington, NC, USA
[7] The Department of Marine Science, The University of North Carolina at Chapel Hill, Chapel Hill, North Carolina, USA


### **Supplemental Figures**

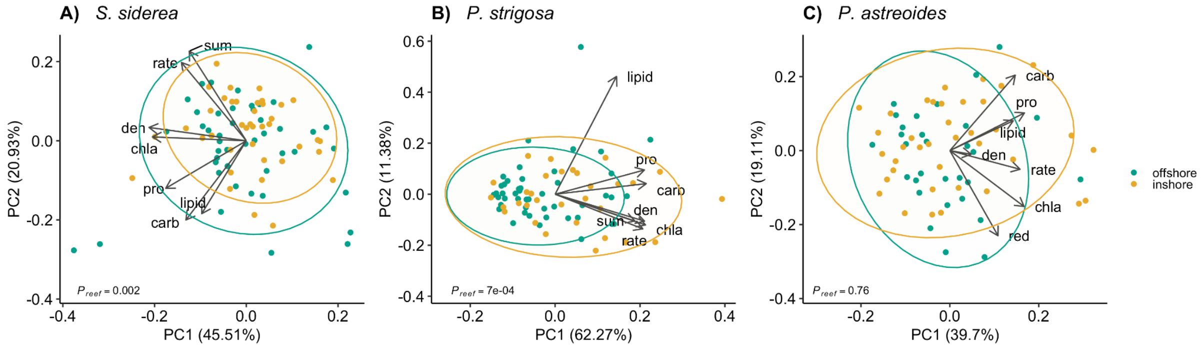


**Figure S1**. Principal component analysis (PCA) of all coral holobiont physiological parameters for (**A**) *S. siderea*, (**B**) *P. strigosa*, and (**C**) *P. astreoides* depicted by natal reef environment (offshore green; inshore yellow). Arrows represent significant ($p < 0.05$) correlation vectors for physiological parameters and ellipses represent 95% confidence based on multivariate t-distributions.

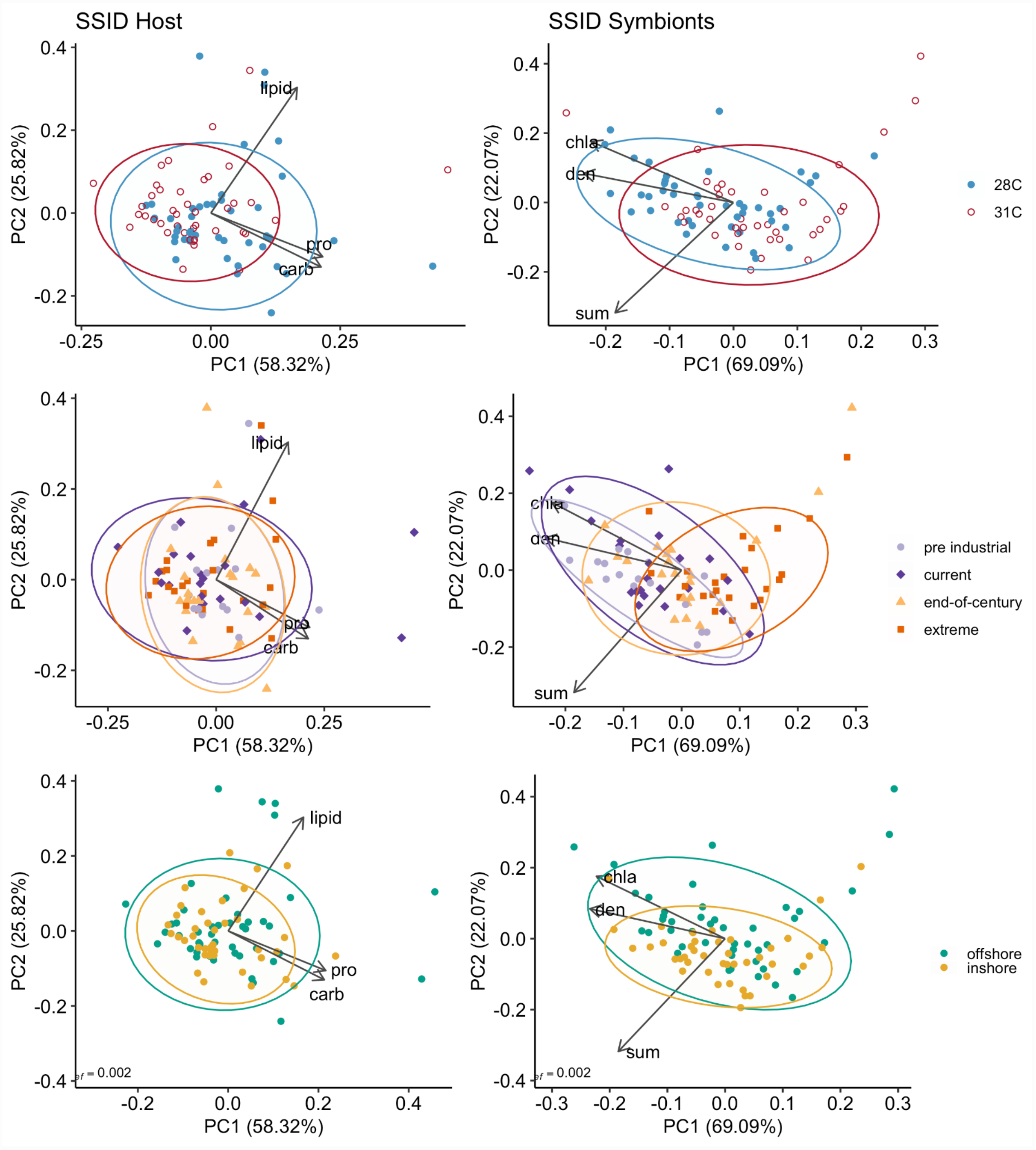


**Figure S2.** Principal component analysis (PCA) of *S. siderea* coral host (protein, lipid, carbohydrate; left) or algal symbiont (chlorophyll a, symbiont density, colour intensity; right) physiological parameters by temperature (28 °C blue; 31 °C red), $pCO_2$ (pre industrial [300 µatm], light purple; current day [420 µatm], dark purple; end-of-century [680 µatm], light orange; extreme [3290 µatm], dark orange), and natal reef environment (offshore green; inshore yellow). Arrows represent significant ($p < 0.05$) correlation vectors for physiological parameters and ellipses represent 95% confidence based on multivariate t-distributions.

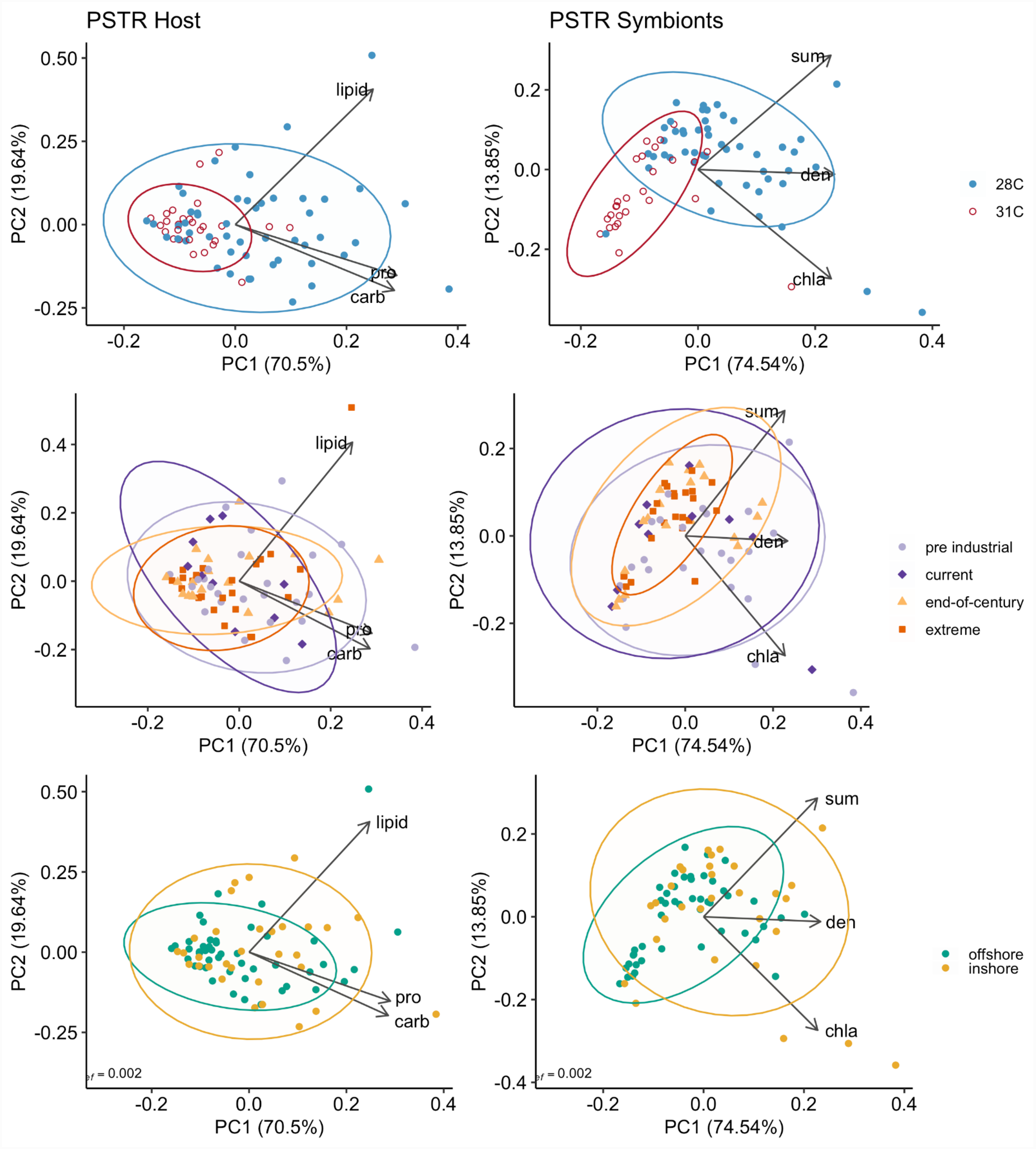


**Figure S3.** Principal component analysis (PCA) of *P. strigosa* coral host (protein, lipid, carbohydrate; left) or algal symbiont (chlorophyll a, symbiont density, colour intensity; right) physiological parameters by temperature (28 °C blue; 31 °C red), $pCO_2$ (pre industrial [300 µatm], light purple; current day [420 µatm], dark purple; end-of-century [680 µatm], light orange; extreme [3290 µatm], dark orange), and natal reef environment (offshore green; inshore yellow). Arrows represent significant ($p < 0.05$) correlation vectors for physiological parameters and ellipses represent 95% confidence based on multivariate t-distributions.

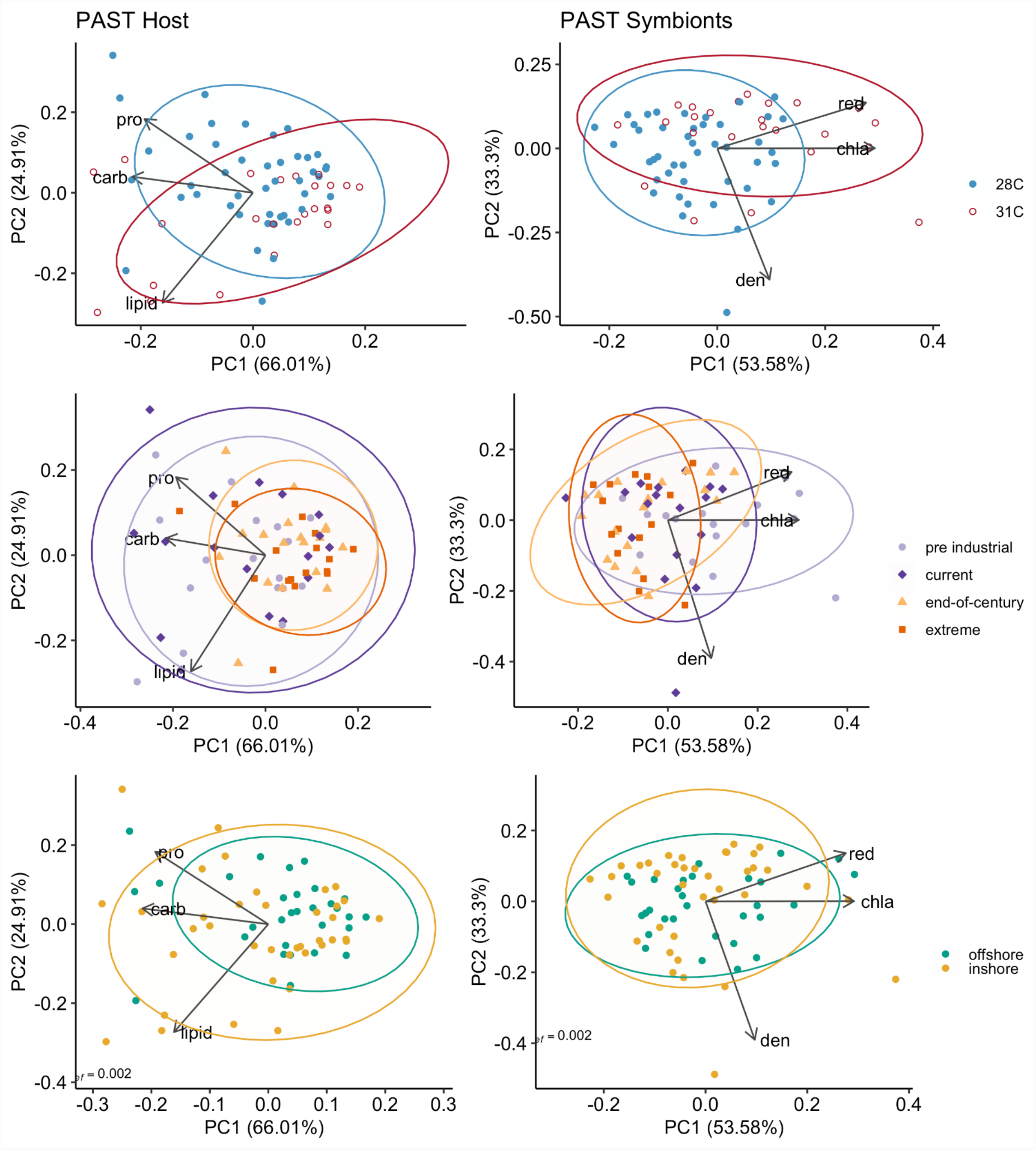


**Figure S4.** Principal component analysis (PCA) of *P. asteroides* coral host (protein, lipid, carbohydrate; left) or algal symbiont (chlorophyll a, symbiont density, colour intensity; right) physiological parameters by temperature (28 °C blue; 31 °C red), $pCO_2$ (pre industrial [300 µatm], light purple; current day [420 µatm], dark purple; end-of-century [680 µatm], light orange; extreme [3290 µatm], dark orange), and natal reef environment (offshore green; inshore yellow). Arrows represent significant ($p < 0.05$) correlation vectors for physiological parameters and ellipses represent 95% confidence based on multivariate t-distributions.

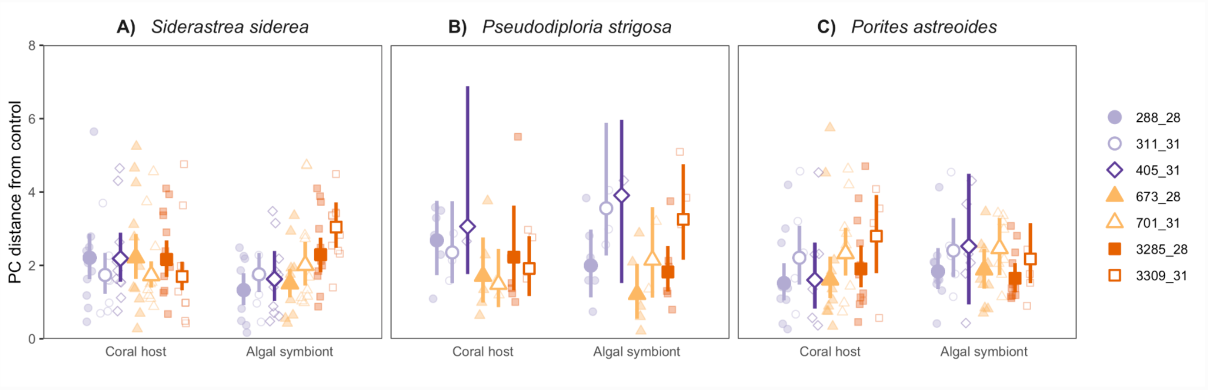


**Figure S5.** Coral host vs. algal symbiont physiological plasticity of (**A**) *S. siderea*, (**B**) *P. strigosa*, and (**C**) *P. astreoides* after 93-day exposure to experimental treatments. Higher values represent greater plasticity in coral holobiont samples. $p$$CO_2$ treatment is depicted by color and shape (pre industrial [300 µatm], light purple; current day [420 µatm], dark purple; end-of-century [680 µatm], light orange; extreme [3290 µatm], dark orange) and temperature is represented as either closed (28 °C) or open (31 °C) symbols. Symbols and bars indicate modeled means and 95% confidence intervals.

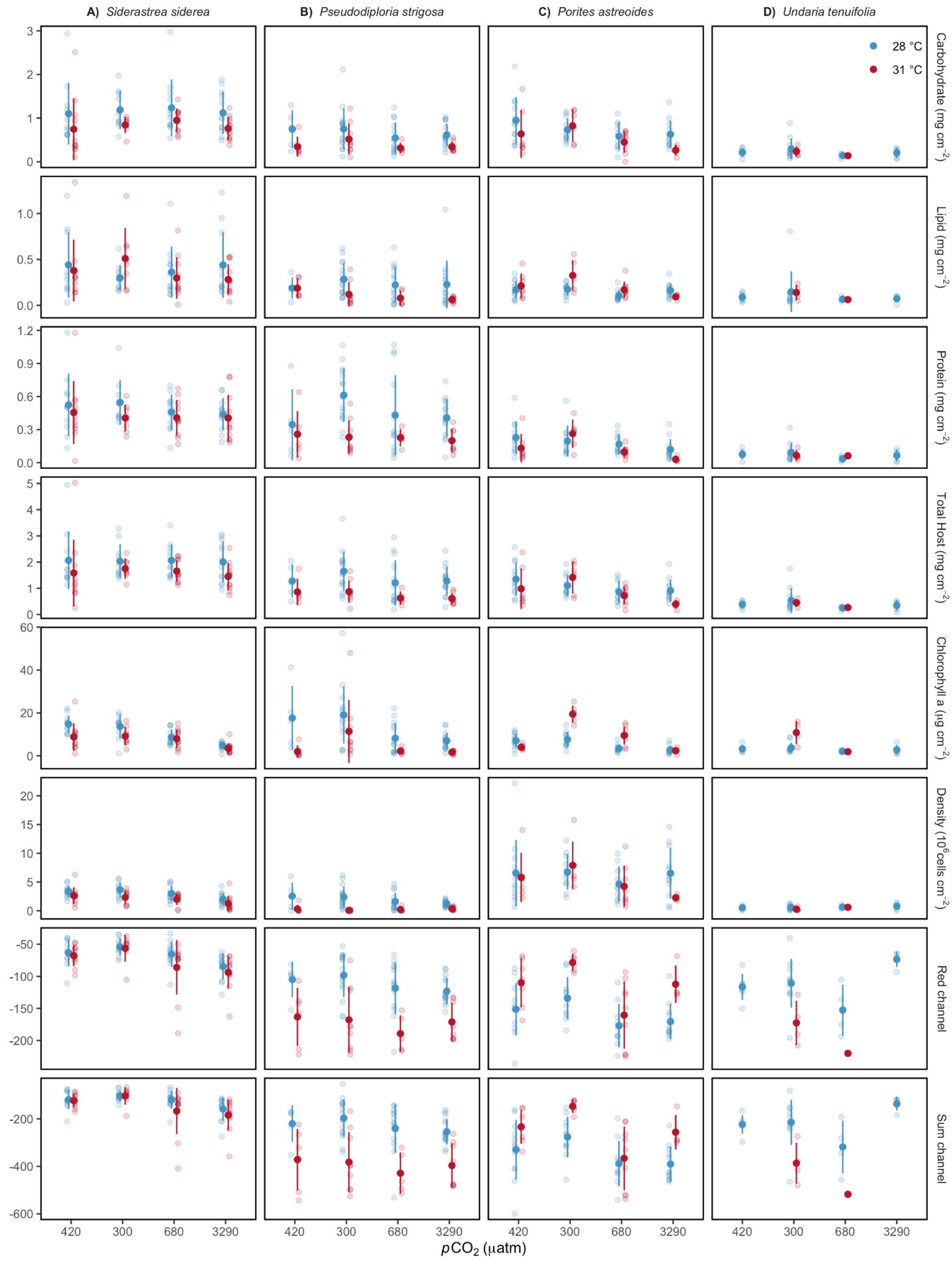


**Figure S6.** Mean (±SE) physiological parameter (each row) measured for (**A**) *S. siderea*, (**B**) *P. strigosa*, (**C**) *P. astreoides*, and (**D**) *U. tenuifolia** at the completion of the 93-day experimental period. *p*$CO_2$ treatment is represented along the x axis and the temperature is depicted by color (28 °C blue; 31 °C red). **U. tenuifolia was included in the original common garden experiment presented in Bove et al 2019 but was not discussed in this manuscript due to high mortality leaving few samples for physiological analyses.*

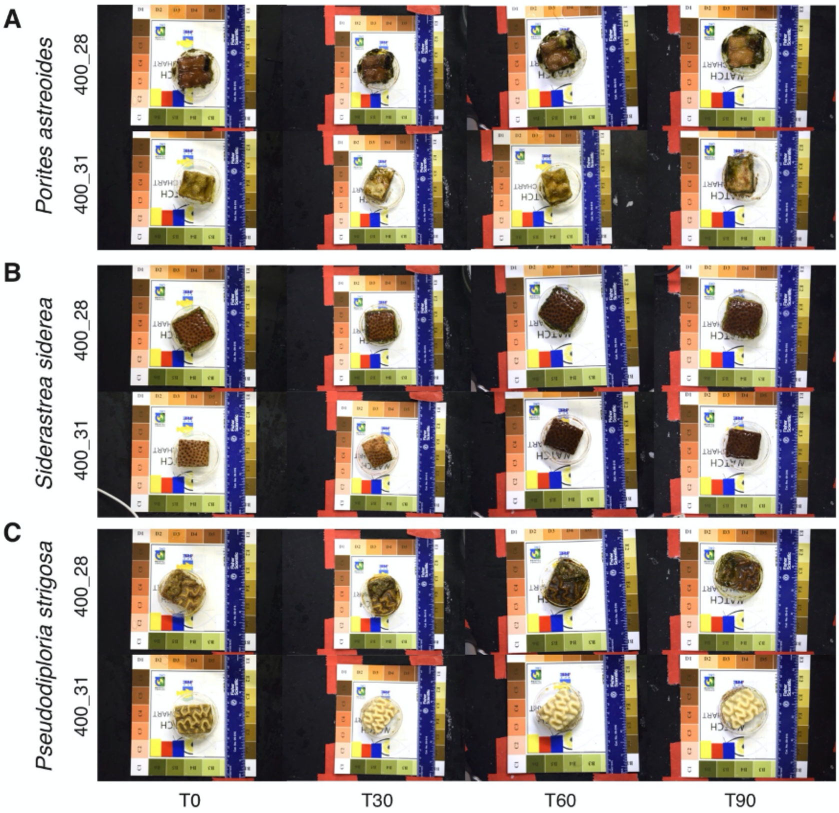


**Figure S7.** Coral colour changes over the experimental period. Representative images of fragments of (**A**) *P. astreoides*, (**B**) *S. siderea*, and (**C**) *P. strigosa* from the same colonies demonstrating change in coral colour over time in either control (420 µatm; 28 °C) or warming (420 µatm; 31 °C) treatments from the start of the experiment (T0) to the end (T90).

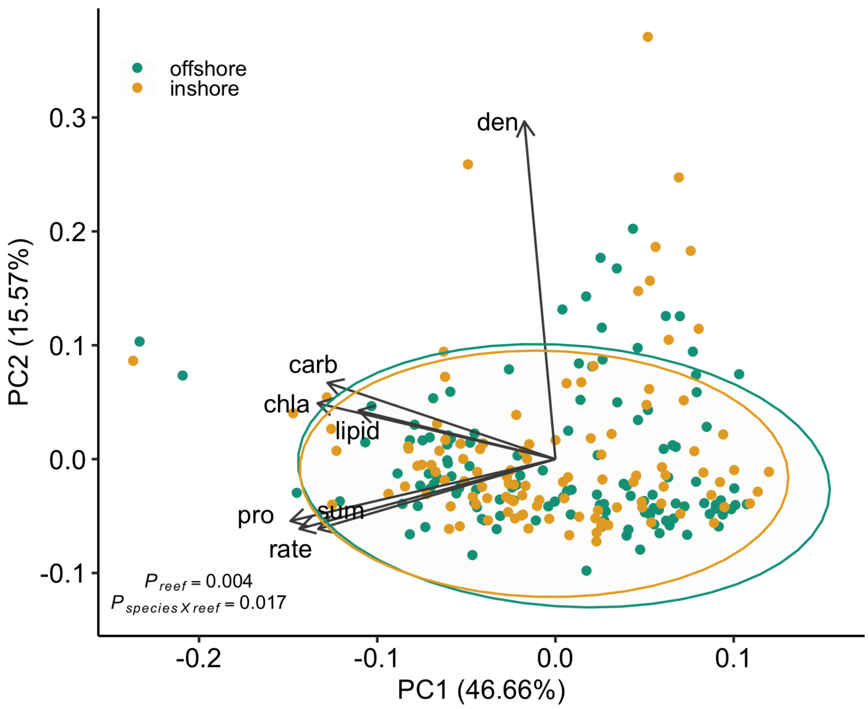


**Figure S8**. Principal component analysis (PCA) comparing the coral holobiont of all three species at the end of the experiment depicted reef environment. Arrows represent significant ($p < 0.05$) correlation vectors for physiological parameters and ellipses represent 95% confidence based on multivariate t-distributions.

### Supplemental Tables

**Table S1.** Model performance comparisons of generalized linear mixed effects models (GLMM) for plasticity assessments to select the best-fit model per species using the package *performance* (version 0.7.0). Akaike information criterion (AIC) was used to select the best-fit model per species. The performance score computes indices of model performance for all models per species at once for comparison across models. The models highlighted in grey were used for bootstrapping estimates and 95% confidence intervals.

| Species | Model formula | AIC | BIC | Conditional $R^2$ | Marginal $R^2$ |
|---|---|---|---|---|---|
| *S. siderea* | reef environment * $pCO_2$ + temperature + (1 \| colony) | 218.8 | 243.5 | 0.506 | 0.322 |
| | reef environment * $pCO_2$* temperature + (1 \| colony) | 223.2 | 259.2 | 0.545 | 0.366 |
| | reef environment * ( $pCO_2$+ temperature) + (1 \| colony) | 220.1 | 247.1 | 0.512 | 0.329 |
| | reef environment + $pCO_2$+ temperature + (1 \| colony) | 221.6 | 239.6 | 0.442 | 0.254 |
| | $pCO_2$+ temperature + (1 \| colony) | 222.1 | 237.8 | 0.37 | 0.088 |
| | reef environment + $pCO_2$* temperature + (1 \| colony) | 225.6 | 248.1 | 0.442 | 0.253 |
| *P. strigosa* | reef environment * $pCO_2$* temperature + (1 \| colony) | 105.6 | 121.5 | 0.397 | 0.317 |
| | reef environment * $pCO_2$+ temperature + (1 \| colony) | 100.8 | 111.8 | 0.309 | 0.261 |
| | reef environment + $pCO_2$* temperature + (1 \| colony) | 101.6 | 112.6 | 0.278 | 0.238 |
| | $pCO_2$+ temperature + (1 \| colony) | 97.5 | 104.8 | 0.232 | 0.188 |
| *P. astreoides* | reef environment * $pCO_2$+ temperature + (1 \| colony) | 145.9 | 167.8 | 0.521 | 0.195 |
| | reef environment * ( $pCO_2$+ temperature) + (1 \| colony) | 147.9 | 171.8 | 0.522 | 0.195 |
| | reef environment * $pCO_2$* temperature + (1 \| colony) | 153.1 | 182.9 | 0.527 | 0.199 |
| | reef environment + $pCO_2$+ temperature + (1 \| colony) | 142.3 | 158.2 | 0.499 | 0.174 |
| | $pCO_2$+ temperature + (1 \| colony) | 140.4 | 154.4 | 0.485 | 0.147 |
| | reef environment + $pCO_2$* temperature + (1 \| colony) | 146.2 | 166.1 | 0.5 | 0.174 |

**Table S2.** PERMANOVA model output from each species using the *adonis2* function with 1500 iterations.

| Species | | Df | Sum of Squares | $R^2$ | F | P-value |
|---|---|---|---|---|---|---|
| | $p CO_2$ | 3 | 61072 | 0.208 | 8.15 | 0.00067 |
| | temperature | 1 | 7471 | 0.025 | 2.99 | 0.09127 |
| *S. siderea* | reef environment | 1 | 24705 | 0.084 | 9.89 | 0.00133 |
| | *Residual* | 80 | 199740 | 0.682 | | |
| | *Total* | 85 | 292988 | 1 | | |
| | reef environment | 1 | 97850 | 0.087 | 14.29 | 0.00067 |
| | $p CO_2$ | 1 | 515604 | 0.456 | 75.29 | 0.00067 |
| *P. strigosa* | temperature | 3 | 30444 | 0.027 | 1.48 | 0.23651 |
| | *Residual* | 71 | 486202 | 0.43 | | |
| | *Total* | 76 | 1130099 | 1 | | |
| | reef environment | 1 | 157 | 0.001 | 0.11 | 0.74151 |
| | temperature | 1 | 25414 | 0.18 | 18.47 | 0.00067 |
| *P. astreoides* | $p CO_2$ | 3 | 30537 | 0.216 | 7.4 | 0.00067 |
| | *Residual* | 62 | 85309 | 0.603 | | |
| | *Total* | 67 | 141417 | 1 | | |

**Table S3.** GLMM output from plasticity assessments for each species. The intercept of each model was set as 300 µatm, 28 °C, and inshore reef environment.

| Species | | Estimate | Standard error | Statistic | P-value |
|---|---|---|---|---|---|
| | *(Intercept)* | 1.085 | 0.172 | 6.3 | 0 |
| | reef environment (offshore) | -0.058 | 0.248 | -0.24 | 0.814 |
| | *p*$CO_2$-current | 0.335 | 0.173 | 1.94 | 0.053 |
| | *p*$CO_2$-EOC | 0.21 | 0.131 | 1.6 | 0.109 |
| | *p*$CO_2$-extreme | 0.419 | 0.132 | 3.17 | 0.002 |
| *S. siderea* | temperature (31°C) | 0.003 | 0.076 | 0.04 | 0.967 |
| | reef environment (offshore):*p*$CO_2$-current | -0.704 | 0.239 | -2.95 | 0.003 |
| | reef environment (offshore):*p*$CO_2$-EOC | -0.409 | 0.197 | -2.08 | 0.037 |
| | reef environment (offshore):*p*$CO_2$-extreme | -0.278 | 0.191 | -1.45 | 0.146 |
| | *Conditional $R^2$* | *0.506* | | | |
| | *Marginal $R^2$* | *0.322* | | | |
| | *(Intercept)* | 1.279 | 0.148 | 8.66 | 0 |
| | *p*$CO_2$-EOC | -0.338 | 0.193 | -1.75 | 0.08 |
| *P. strigosa* | *p*$CO_2$-extreme | -0.059 | 0.187 | -0.31 | 0.753 |
| | temperature (31°C) | 0.227 | 0.173 | 1.31 | 0.19 |
| | *Conditional $R^2$* | *0.232* | | | |
| | *Marginal $R^2$* | *0.188* | | | |
| | *(Intercept)* | 1.038 | 0.121 | 8.61 | 0 |
| | *p*$CO_2$-current | -0.047 | 0.11 | -0.43 | 0.67 |
| | *p*$CO_2$-EOC | 0.032 | 0.075 | 0.44 | 0.664 |
| *P. astreoides* | *p*$CO_2$-extreme | 0.122 | 0.078 | 1.57 | 0.116 |
| | temperature (31°C) | 0.264 | 0.065 | 4.07 | 0 |
| | *Conditional $R^2$* | *0.485* | | | |
| | *Marginal $R^2$* | *0.147* | | | |

**Table S4.** PERMANOVA model output across species using the *adonis2* function with 1500 iterations.

| | Df | Sum of Squares | $R^2$ | F | P-value |
|---|---|---|---|---|---|
| $pCO_2$ | 3 | 149393 | 0.04 | 8.24 | 0.0007 |
| temperature | 1 | 17313 | 0 | 2.87 | 0.0933 |
| reef environment | 1 | 58058 | 0.02 | 9.61 | 0.0047 |
| species | 2 | 1642613 | 0.42 | 135.9 | 0.0007 |
| temperature:species | 2 | 553351 | 0.14 | 45.78 | 0.0007 |
| $pCO_2$:species | 6 | 90865 | 0.02 | 2.51 | 0.024 |
| reef environment:species | 2 | 77259 | 0.02 | 6.39 | 0.004 |
| *Residual* | *214* | *1293204* | *0.33* | | |
| *Total* | *231* | *3882055* | *1* | | |

**Table S5.** PERMANOVA model output of coral host or algal symbiont physiology per species using the *adonis2* function with 1500 iterations depicted in **Figures S2-S4**.

| | Coral host | | | | | Algal symbiont | | | | |
|---|---|---|---|---|---|---|---|---|---|---|
| | **Df** | **Sum of Squares** | **$R^2$** | **F** | **P-value** | **Df** | **Sum of Squares** | **$R^2$** | **F** | **P-value** |
| ***S. siderea*** | | | | | | | | | | |
| $pCO_2$ | 3 | 1 | 0.019 | 0.55 | 0.74883 | 3 | 61056 | 0.208 | 8.15 | 0.00067 |
| temperature | 1 | 3 | 0.075 | 6.65 | 0.00333 | 1 | 7468 | 0.025 | 2.99 | 0.10127 |
| reef environment | 1 | 0 | 0.006 | 0.52 | 0.57295 | 1 | 24705 | 0.084 | 9.9 | 0.00266 |
| *Residual* | *80* | *30* | *0.901* | | | *80* | *199684* | *0.682* | | |
| *Total* | *85* | *34* | *1* | | | *85* | *292913* | *1* | | |
| ***P. strigosa*** | | | | | | | | | | |
| $pCO_2$ | 3 | 1 | 0.041 | 1.23 | 0.3058 | 3 | 26899 | 0.024 | 1.31 | 0.28181 |
| temperature | 1 | 3 | 0.147 | 13.12 | 0.00067 | 1 | 515173 | 0.456 | 75.24 | 0.00067 |
| reef environment | 1 | 0 | 0.02 | 1.75 | 0.15656 | 1 | 101793 | 0.09 | 14.87 | 0.00067 |
| *Residual* | *71* | *14* | *0.793* | | | *71* | *486140* | *0.43* | | |
| *Total* | *76* | *18* | *1* | | | *76* | *1130005* | *1* | | |
| ***P. astreoides*** | | | | | | | | | | |
| $pCO_2$ | 3 | 2 | 0.136 | 3.48 | 0.01532 | 3 | 29037 | 0.205 | 7.04 | 0.00133 |
| temperature | 1 | 0 | 0.036 | 2.76 | 0.08328 | 1 | 26338 | 0.186 | 19.15 | 0.00067 |
| reef environment | 1 | 0 | 0.021 | 1.64 | 0.18121 | 1 | 724 | 0.005 | 0.53 | 0.47768 |
| *Residual* | *62* | *10* | *0.807* | | | *62* | *85288* | *0.603* | | |
| *Total* | *67* | *13* | *1* | | | *67* | *141387* | *1* | | |

**Table S6.** Model performance comparisons of generalized linear mixed effects models (GLMM) for plasticity assessments to select the best-fit model comparing host and symbiont using the package *performance* (version 0.7.0). Akaike information criterion (AIC) was used to select the best-fit model. The performance score computes indices of model performance for all models per species at once for comparison across models. The models highlighted in grey were used for bootstrapping estimates and 95% confidence intervals.

| Model formula | AIC | BIC | Conditional $R^2$ | Marginal $R^2$ |
|---|---|---|---|---|
| species * part * reef environment * $pCO_2$ * temperature + (1 \| colony) | 923.1 | 1219.4 | 0.43 | 0.318 |
| species * part * reef environment * ($pCO_2$ + temperature) + (1 \| colony) | 902.1 | 1124.3 | 0.391 | 0.279 |
| reef environment * species * part * ($pCO_2$ + temperature) + (1 \| colony) | 902.1 | 1124.3 | 0.391 | 0.279 |
| reef environment * species * part * ($pCO_2$ + temperature) + (1 \| colony) | 902.1 | 1124.3 | 0.391 | 0.279 |
| species * part * ($pCO_2$ + temperature + reef environment) + (1 \| colony) | 883.9 | 1024.7 | 0.323 | 0.213 |
| species * part * ($pCO_2$ + temperature) + (1 \| colony) | 882 | 1000.5 | 0.28 | 0.139 |
| species * part * ($pCO_2$ + temperature) + reef environment + (1 \| colony) | 883.5 | 1005.7 | 0.282 | 0.144 |
| species * part * reef environment * $pCO_2$ + temperature + (1 \| colony) | 904.6 | 1086.1 | 0.338 | 0.225 |
| species * part * $pCO_2$ + temperature + (1 \| colony) | 890.2 | 990.2 | 0.235 | 0.093 |
| species * reef environment + part + $pCO_2$ + temperature + (1 \| colony) | 881.2 | 929.4 | 0.205 | 0.095 |
| species * part * reef environment + $pCO_2$ * temperature + (1 \| colony) | 890.2 | 964.2 | 0.222 | 0.113 |

**Table S7.** GLMM output from plasticity assessments for coral host vs. algal symbiont for each species. The intercept of each model was set as coral host, *S. siderea*, 300 µatm, and 28 °C.

| | Estimate | Standard error | Statistic | P-value |
|---|---|---|---|---|
| *(Intercept)* | 0.8 | 0.172 | 4.66 | 0 |
| PSTR | 0.2 | 0.29 | 0.69 | 0.491 |
| PAST | -0.357 | 0.244 | -1.47 | 0.143 |
| symbionts | -0.515 | 0.185 | -2.78 | 0.005 |
| $p\mathrm{CO_2}$-current | 0.235 | 0.214 | 1.1 | 0.272 |
| $p\mathrm{CO_2}$-EOC | 0.006 | 0.164 | 0.03 | 0.972 |
| $p\mathrm{CO_2}$-extreme | -0.021 | 0.161 | -0.13 | 0.896 |
| temperature (31°C) | -0.252 | 0.128 | -1.97 | 0.049 |
| PSTR:symbionts | 0.222 | 0.308 | 0.72 | 0.472 |
| PAST:symbionts | 0.685 | 0.27 | 2.54 | 0.011 |
| PSTR:$p\mathrm{CO_2}$-current | -0.088 | 0.592 | -0.15 | 0.882 |
| PAST:$p\mathrm{CO_2}$-current | -0.569 | 0.339 | -1.68 | 0.093 |
| PSTR:$p\mathrm{CO_2}$-EOC | -0.451 | 0.298 | -1.51 | 0.131 |
| PAST:$p\mathrm{CO_2}$-EOC | 0.05 | 0.24 | 0.21 | 0.835 |
| PSTR:$p\mathrm{CO_2}$-extreme | -0.154 | 0.288 | -0.54 | 0.592 |
| PAST:$p\mathrm{CO_2}$-extreme | 0.236 | 0.245 | 0.96 | 0.336 |
| PSTR:temperature (31°C) | 0.071 | 0.25 | 0.29 | 0.775 |
| PAST:temperature (31°C) | 0.608 | 0.2 | 3.04 | 0.002 |
| symbionts:$p\mathrm{CO_2}$-current | -0.299 | 0.298 | -1 | 0.316 |
| symbionts:$p\mathrm{CO_2}$-EOC | 0.134 | 0.227 | 0.59 | 0.556 |
| symbionts:$p\mathrm{CO_2}$-extreme | 0.564 | 0.226 | 2.5 | 0.013 |
| symbionts:temperature (31°C) | 0.524 | 0.181 | 2.89 | 0.004 |
| PSTR:symbionts:$p\mathrm{CO_2}$-current | 0.362 | 0.818 | 0.44 | 0.658 |
| PAST:symbionts:$p\mathrm{CO_2}$-current | 0.673 | 0.473 | 1.42 | 0.155 |
| PSTR:symbionts:$p\mathrm{CO_2}$-EOC | -0.177 | 0.418 | -0.42 | 0.672 |
| PAST:symbionts:$p\mathrm{CO_2}$-EOC | -0.181 | 0.335 | -0.54 | 0.59 |
| PSTR:symbionts:$p\mathrm{CO_2}$-extreme | -0.482 | 0.401 | -1.2 | 0.229 |
| PAST:symbionts:$p\mathrm{CO_2}$-extreme | -0.891 | 0.341 | -2.61 | 0.009 |
| PSTR:symbionts:temperature (31°C) | 0.191 | 0.35 | 0.54 | 0.586 |
| PAST:symbionts:temperature (31°C) | -0.606 | 0.281 | -2.16 | 0.031 |
| *Conditional* $R^2$ | *0.28* | | | |
| *Marginal* $R^2$ | *0.139* | | | |
